# Altered integration of excitatory inputs onto the basal dendrites of layer 5 pyramidal neurons in a mouse model of Fragile X Syndrome

**DOI:** 10.1101/2022.03.14.484306

**Authors:** Diana E. Mitchell, Soledad Miranda-Rottmann, Maxime Blanchard, Roberto Araya

## Abstract

To uncover how synaptic integration of feedforward sensory inputs is affected in autism spectrum disorders (ASD), we used two-photon uncaging of caged glutamate to activate two clustered spines in the basal dendrites of layer 5 (L5) pyramidal neurons from a mouse model of Fragile X syndrome (FXS), the most common genetic cause of ASD. While subthreshold excitatory inputs integrate linearly in wild-type animals, surprisingly those with FXS summate sublinearly, contradicting what would be expected of a hyperexcitable cortex typically associated with ASD. Knockdown of the regulatory β4 subunit of BK channels, rescues the synaptic integration, a result that was corroborated with numerical simulations. Taken together, these findings suggest that there is a differential impairment in the integration of feedforward sensory and feedback predictive inputs in L5 pyramidal neurons in FXS and potentially other forms of ASD. These results challenge the traditional view that FXS and other ASD are characterized by sensory hypersensitivity, but instead by hyposensitivity of sensory inputs and hypersensitivity of predictive inputs onto cortical neurons.

## MAIN

Fragile-X syndrome (FXS) is the most frequent form of inherited intellectual disability and the most common known cause of autism (Jacquemont et al., 2007; Wang et al., 2012). FXS and other neurodevelopmental disorders associated with autism spectrum disorders (ASD) have a negative impact in most, if not all, everyday life activities, with a global prevalence that has been growing steeply since the 1990’s (Rice, 2013). FXS occurs because of the inactivation of the Fragile X Mental Retardation 1 (FMR1) gene, which encodes the FXS Protein (FMRP), a polyribosome-associated RNA-binding protein that inhibits the translation of bound mRNAs, especially at synapses (Darnell et al., 2011; Hale et al., 2021).

At the cellular level, it has been shown that spines (tiny protrusions covering the dendrites of cortical pyramidal and spiny stellate neurons that constitute the postsynaptic element of ~95% of all the excitatory synapses (Gray, 1959; Colonnier, 1968; Arellano et al., 2007)) are abnormally long and dense in mice models of FXS (*Fmr1*-KO mice) (Rudelli et al., 1985; Irwin et al., 2001). These synaptic alterations suggest the presence of defects in the transmission, plasticity and integration of excitatory inputs in FXS (Portera-Cailliau, 2012; He and Portera-Cailliau, 2013). At the circuit level, the neocortex of *Fmr1*-KO mice has been found to be hyperexcitable (Bureau et al., 2008; Gibson et al., 2008; Rotschafer and Razak, 2013; Zhang et al., 2014; Booker et al., 2019) with pyramidal neurons exhibiting abnormally high and synchronous firing, leading to recurrent bursting (Goncalves et al., 2013) – something long assumed to cause sensory hypersensitivity (reviewed in: Liu et al., 2021).

A key function of the neocortex is to associate external sensory information with an internal representation of the world to make predictions about the future (Felleman and Van Essen, 1991; Shipp, 2007; Larkum, 2013). Specifically, layer 5 (L5) pyramidal neurons integrate sensory inputs (feedforward) in their basal dendrites with information from other cortical areas (feedback) at the apical dendrites in layer 1 (Larkum, 2013). Feedback information provides contextual or invariant predictive information built from previous experiences, while feedforward inputs provide external sensory information. Thus, to convert an invariant prediction into a specific prediction about the world (cognition and conscious perception) the brain has to combine feedforward with feedback information (Lamme et al., 1998; Pascual-Leone and Walsh, 2001; Soltani and Koch, 2010; Boly et al., 2011). L5 pyramidal neurons, vertically spanning all cortical layers, are the main candidates believed to perform this task (Larkum, 2013; Takahashi et al., 2016). Hence, defects in the processing and integration of excitatory inputs at the level of single dendritic spines in the basal and/or distal apical tuft dendrites of L5 pyramidal neurons could contribute to the functional cortical defects associated with neurological disorders such as FXS and ASD. Indeed, previous work has shown an increased stimulus-evoked response at the soma of *Fmr1*-KO L5 pyramidal neurons as well as an enhanced temporal summation of excitatory inputs in their distal (apical tuft) dendrites in response to multiple pulses of electrical stimulation in layer 1 (Zhang et al., 2014). It remains unknown; however, how sensory inputs are processed at the level of individual synapses in the dendrites of *Fmr1*-KO L5 pyramidal neurons, and whether the synaptic integration impairments are equally or differentially affected within feedforward and feedback pathways in L5 pyramidal neurons in FXS.

Here, we aim to uncover if the integration of sensory inputs at the level of single synapses in the basal dendrites of L5 pyramidal neurons is altered in *Fmr1*-KO mice and, if so, explore the underlying spine mechanisms responsible for any observed differences. To do so, we first determined how near-simultaneous inputs onto neighboring spines are integrated in L5 pyramidal neuron dendrites from *Fmr1*-KO versus wild-type (WT) mice. As previously demonstrated (Araya et al., 2006a), we found that the basal dendrites of WT L5 pyramidal neurons integrate inputs linearly before the generation of a dendritic spike, whereas, surprisingly, synaptic inputs in L5 pyramidal neuron basal dendrites summate sublinearly in *Fmr1*-KO mice, contradicting what would be expected of a hyperexcitable cortex typically associated with FXS. We reasoned that these integration impairments are due to spine channelopathies (Deng and Klyachko, 2021). Previous work has revealed that FMRP interacts with the β4 regulatory subunit of large-conductance voltage- and calcium-activated potassium (BK) channels (Deng et al., 2013), which are localized to dendritic spines where they play a role contributing to synaptic efficacy in L5 pyramidal neurons (Tazerart et al., 2022). Thus, in WT mice, FMRP can sequester the β4 subunit, as has been shown in the hippocampal pyramidal neurons (Deng et al., 2013), and prevent it from interacting with BK channels, whereas in *Fmr1*-KO animals, an increased binding of BK channels and its β4 subunit would occur. The β4 subunit decreases the probability of BK channels opening at low calcium concentrations and increases it at high calcium concentrations while also slowing down activation and deactivation kinetics (Brenner et al., 2000; Torres et al., 2007), which would ultimately affect L5 pyramidal neuron integrative properties. Hence, we predicted that the sublinear integration in the basal dendrites of *Fmr1*-KO L5 pyramidal neurons was due to altered BK channel activity in the absence of FMRP, compared to WT neurons. Here, we found that knockdown of the β4 subunit of BK channels, using short hairpin RNA (shRNA), rescues the linear integration of subthreshold inputs in the basal dendrites of *Fmr1*-KO L5 pyramidal neurons while injection of a scrambled version has no effect. Indeed, numerical simulations corroborate these experimental results, showing that synaptic integration occurs linearly only when the kinetics of the α-subunit of spine BK channels are modeled, while inputs integrate sublinearly when the β4 subunit are incorporated into the spine. Taken together, the results from these experiments highlight the need to study sensory (feedforward) and contextual/predictive (feedback) pathways independently in FXS and in other forms of ASD and to move away from the idea of that FXS is solely characterized by sensory hypersensitivity, but instead by sensory hyposensitivity and hypersensitivity of predictive inputs. Finally, these findings help to uncover the role of ion channels in excitatory input integration and identify novel and localized dendritic targets for the design of specific treatment options to alleviate symptoms associated with FXS.

## RESULTS

### Sublinear integration in the basal dendrites of L5 pyramidal neurons in *Fmr1*-KO mice

To study synaptic integration in the basal dendrites of L5 pyramidal neurons in acute brain slices of mouse visual cortex from WT and *Fmr1*-KO mice, we applied 2P uncaging of caged glutamate (4-methoxy-7-nitroindolinyl glutamate (MNI)-glutamate, 2.5mM see Material and Methods) at two individual spines separately and then together near-simultaneously (Fig. 1a). Whole-cell patch clamp recordings in current-clamp were performed to measure the uncaging (u)-evoked excitatory postsynaptic potentials (uEPSP) at the soma. This technique allows for the precise activation of individual dendritic spines with responses that are similar to those from physiological activation of single synapses (Matsuzaki et al., 2001; Araya et al., 2006a; Araya et al., 2006b; Fino et al., 2009; Araya et al., 2013; Araya, 2014; Araya et al., 2014; Mitchell et al., 2019; Tazerart et al., 2020; Tazerart et al., 2022). As shown previously (Araya et al., 2006a), subthreshold synaptic inputs onto spines in the basal dendrites of WT L5 pyramidal neurons integrate linearly (Fig. 1a). Specifically, we found that activating individual spines triggered uEPSPs (black traces in upper panel of Fig. 1a), which added linearly to accurately predict the response when two spines were activated near-simultaneously (compare black and blue dashed traces in upper right panel of Fig. 1a). We quantified these results by calculating linearity indices, using either the peak or integral of the uEPSP as well as a gain measure (see Methods). Linearity indices were not significantly different from 100% for WT L5 pyramidal neurons when we considered each experiment individually (100.45 ± 1.74%, P = 0.78, for peak uEPSP, and 102.82 ± 2.15% P = 0.20 for uEPSP integral, Wilcoxon test, n = 40 spine pairs; Fig. 1b) or when we averaged each individual experiment per mouse (98.35 ± 2.82%, P = 0.49, Wilcoxon test, n = 10 mice for peak uEPSP, and 99.90 ± 3.79% P = 0.32, Wilcoxon test, n = 10 mice for uEPSP integral; Fig. 1c). We next computed a gain measure, which compares the time-varying uEPSP in response to the activation of 2 spines to that predicted from the linear summation of individual responses, with a measure of one being a perfect match. We found that the gain measure was not significantly differently from one for WT L5 pyramidal neurons (*individual experiments*: 1.03 ± 0.02, P = 0.29, n = 40 spine pairs; *average per mouse*: 1.00 ± 0.02, P =0.98, n = 10 mice; Wilcoxon test, Fig. 1b-c). Surprisingly, we observed uEPSP responses that were smaller in amplitude and integral than expected based on the sum of individual responses in *Fmr1*-KO mice (compare red and blue dashed traces in lower right panel of Fig. 1a) with linearity indices significantly lower than 100% (*individual experiments*: 91.27 ± 1.47%, P < 0.0001 for peak uEPSP and 89.58 ± 1.90%, P < 0.0001 for uEPSP integral, Wilcoxon test, n = 74 spine pairs; *average per mice*: 88.56 ± 1.79%, P < 0.0001 for peak uEPSP and 88.28 ± 1.89 %, P < 0.0001 for uEPSP integral, Wilcoxon test, n = 16 mice; Fig. 1b-c) and the gain measure significantly lower than one (*individual experiments*: 0.89 ± 0.02, P < 0.0001, n = 74 spine pairs, Wilcoxon test; *average per mice*: 0.87 ± 0.03, P < 0.001, n = 16 mice, Wilcoxon test; Fig. 1b-c). These results were surprising since they contradict what would be expected from a hyperexcitable cortex, that has been described in FXS.

**Figure 1:**
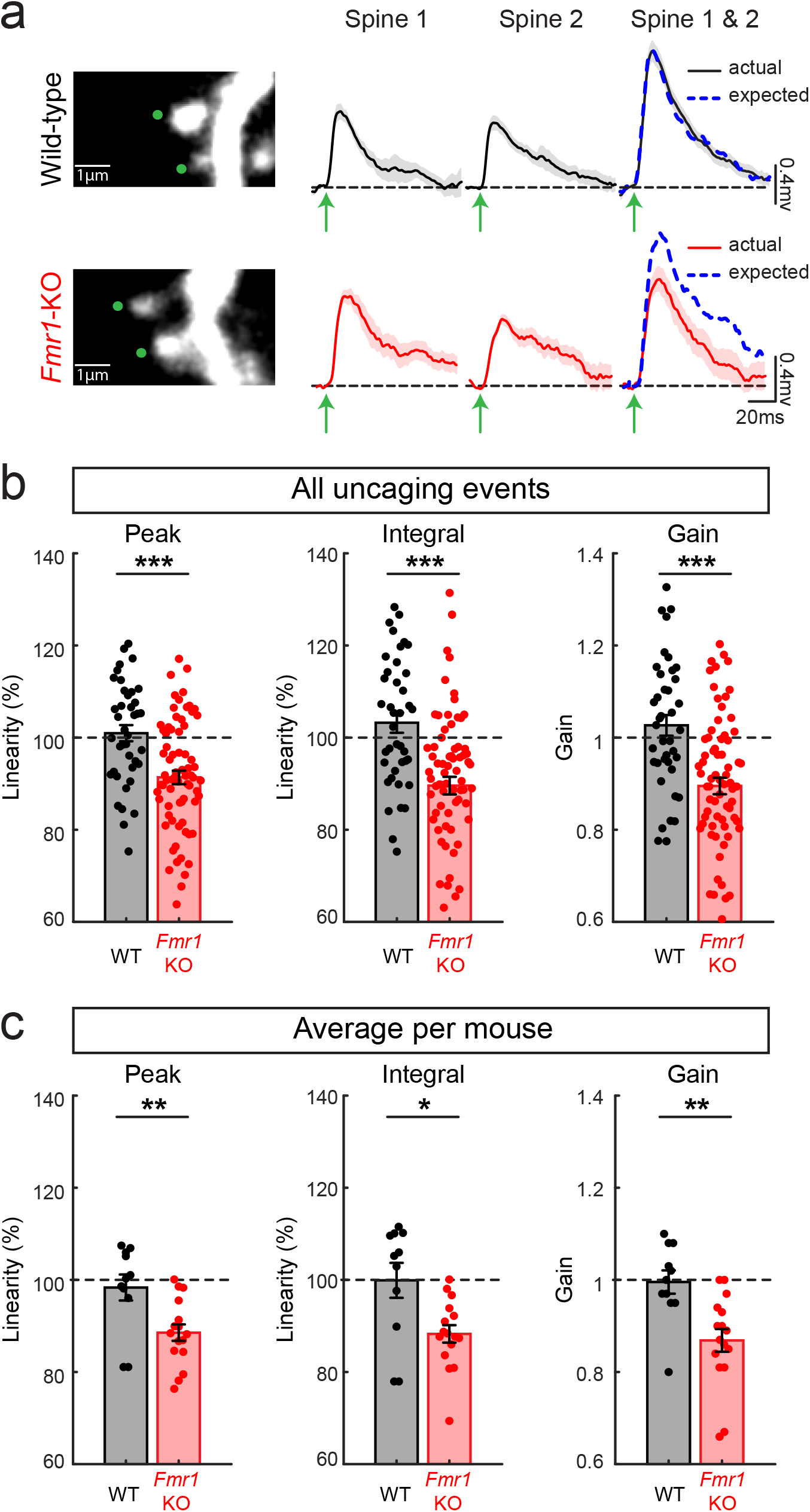
Sublinear summation of excitatory synaptic inputs in basal dendrites of Fmr1-KO L5 pyramidal neurons. (**a**) Representative proximal dendrite selected for 2P glutamate uncaging to activate spines from WT (top panels) and *Fmr1*-KO L5 pyramidal neurons (bottom panels). Green dots indicate the site for uncaging. Spines were first activated individually (Spine 1 or Spine 2) and then together (Spine 1 & Spine 2). Dashed blue traces correspond to the linear sum of individual events of each spine. (**b**) Observed response has smaller amplitude, integral and gain than expected based on the sum of individual responses in *Fmr1*-KO mice, while in WT mice, the responses summate linearly. (**c**) As in **b**, but values were averaged per mouse.

We found no difference in the intrinsic cellular properties between *Fmr1*-KO versus WT L5 pyramidal neurons (Supplementary Fig. 1). When we activated one (individual) versus two (combined) spines, we found that individual and combined uEPSP responses were smaller in the basal dendrites of *Fmr1*-KO versus WT L5 pyramidal neurons (Supplementary Fig. 2). This effect was washed out, however, when experiments were pooled per mouse (Supplementary Fig. 2). Thus, in some animals, there was an overrepresentation of experiments yielding smaller uEPSP sizes. The morphology of activated spines was comparable for *Fmr1*-KO and WT mice (Supplementary Fig. 2).

### Expression of BK channel and its β4-subunit in neocortex of WT versus *Fmr1*-KO mice

We next sought to identify the mechanism for this sublinear integration of spine activation in the basal dendrites of *Fmr1*-KO L5 pyramidal neurons. Large conductance calcium-activated potassium (BK) channels are homogeneously distributed along the dendritic tree of L5 pyramidal neurons and are believed to play an important role in synaptic transmission and integration (Bock and Stuart, 2016; Tazerart et al., 2022). BK channels are high-conductance K^+^ channel formed by a tetramer of α subunits. Membrane depolarization and intracellular Ca^2+^ activate these channels, and in many tissues auxiliary subunits modulate their kinetics (reviewed in: Latorre et al., 2017). For example, it has been demonstrated in hippocampal CA3 pyramidal neurons that FMRP modulates BK channel activity by sequestering its regulatory β4 subunit (Deng et al., 2013). The β4 subunit decreases the probability of BK channel openings at low calcium concentrations but increases the probability of channel openings at high calcium concentrations while also slowing down activation and deactivation kinetics (Brenner et al., 2000; Torres et al., 2007). We thus predicted that *Fmr1*-KO L5 pyramidal neurons exhibit altered BK channel activity in the absence of FMRP, compared to WT neurons, which would ultimately affect L5 pyramidal neuron integrative properties. Using Western blot analysis of total protein (TP) and synaptoneurosomes (SN) prepared from visual cortex, we show that αBK subunits are enriched in synapses and there is no expression difference between WT and *Fmr1*-KO mice (WT: 1.00 ± 0.13 in SN vs 7.31 ± 0.31 in TP; KO: 1.54 ± 0.09 in SN vs 7.31±0.44 in TP, n=3, p<0.001) (Fig. 2a-b). By contrast, β4 subunits are not enriched in synapses in WT mice but are enriched in those of *Fmr1*-KO mice (WT: 0.90 ± 0.19 in SN vs 0.51 ± 0.09 in TP; *Fmr1*-KO: 1.04 ± 0.36 in SN vs 2.26 ±0.17 in TP, n=3 mixed, p<0.001), suggesting that the absence of FMRP to sequester β4 subunits from its interaction with BK channels in spines increases β4 immunoreactivity in the synapse (Fig. 2c-d). In addition, immunofluorescence data in transgenic mice expressing GFP in L5 pyramidal neurons shows that αBK (Fig. 2e) and β4 (Fig. 2f) subunits are present in basal dendrites and spines from WT and *Fmr1*-KO L5 pyramidal neurons. Interestingly, the images suggest an increased β4 immunoreactivity in dendrites and dendritic spines of *Fmr1*-KO compared to WT. Moreover, we have previously shown using immunoelectron microscopy, that αBK subunits are localized to dendritic spines in the basal dendrites from L5 pyramidal neurons with nanoscale resolution (Tazerart et al., 2022). Taken together, these experiments along with previous findings provide evidence that we predict would alter BK channel activity in spines from *Fmr1*-KO L5 pyramidal neurons thus affecting integration of synaptic inputs in these neurons.

**Figure 2:**
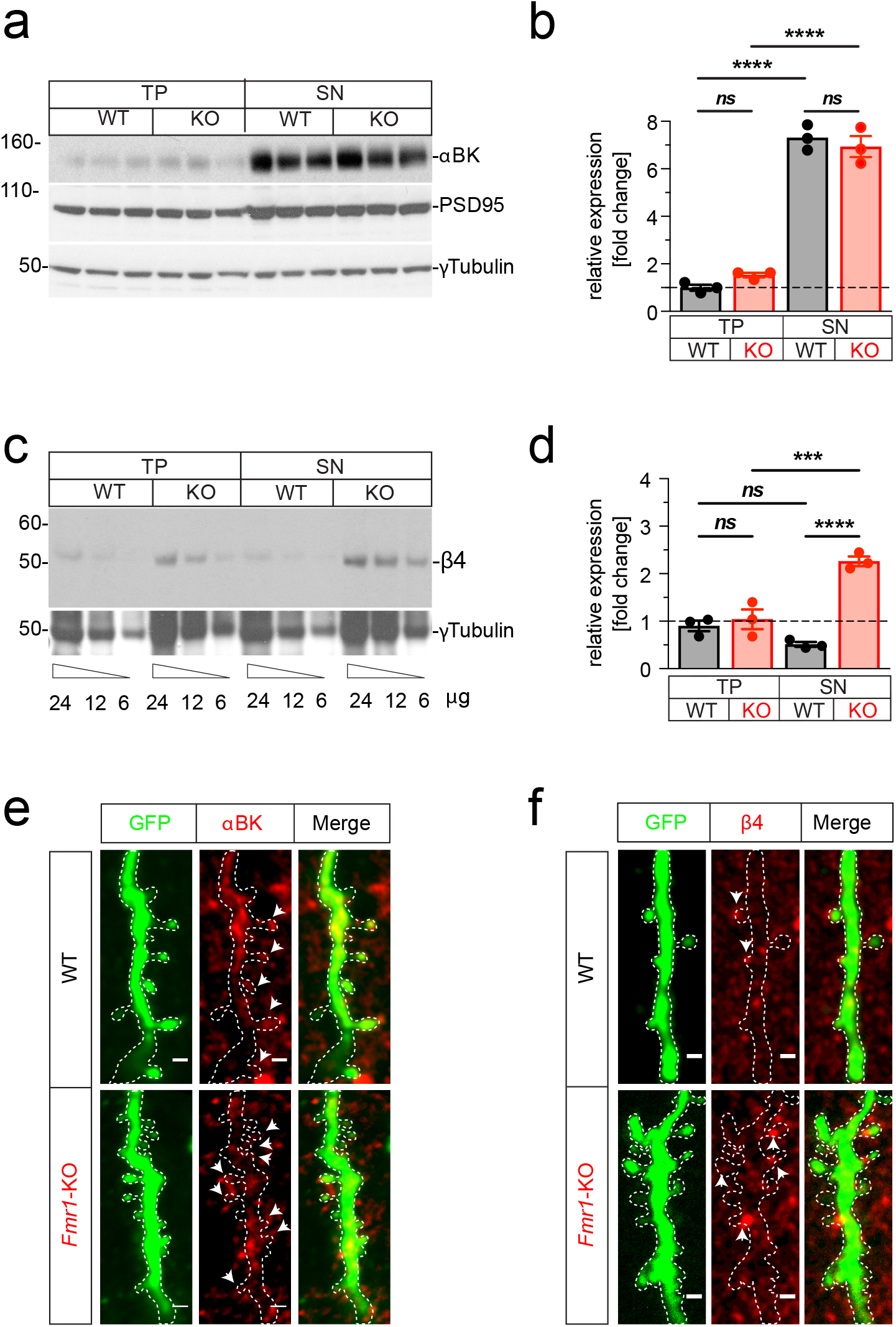
Expression of αBK and β4 subunits in synapses of *Fmr1*-KO mice. (**a-b**) Mouse visual cortex total protein (TP) and Synaptoneurosome (SN) fractions were analysed by Western Blot with specific monoclonal antibodies and normalized by the expression of γ-tubulin. Expression of αBK (20 μg/line) (**a**). Synaptic enrichment in the SN fraction was confirmed by the expression of the post-synaptic density (PSD) marker PSD95. αBK expression normalized by γ-tubulin and by the expression in WT/TP (n=3, P<0.0001) (**b**). (**c**) Expression of β4. A mix of the same samples was loaded at decreasing concentrations (24, 12, 6 μg/line). (**d**) β4 expression normalized by γ-tubulin and by the expression in WT/TP (n=3, P<0.05, P<0.0001). (**e-f**) Expression of channel subunits in excitatory synapses was studied by immunofluorescent detection of αBK (**e**) and β4 (red) (**f**) in transgenic mice expressing GFP in L5 pyramidal neuron (green). Dendritic spine channel expression is shown in one <1 μm confocal optical slice. Arrow heads indicate spines with detectable expression of the BK subunit. Scale bar 1 μm.

### Knock-down of β4 subunit of BK channels in neocortex of *Frm1*-KO mice

In order to test the hypothesis that the lack of FRMP in *Frm1-KO* mice leads to an elevated binding of the β4 subunit to BK channels in spines and dendrites, thus disrupting the linearity of incoming synaptic inputs, we performed a knockdown of the β4 subunit of BK channels, using short hairpin RNA (shRNA) in *Fmr1*-KO L5 pyramidal neurons (Fig. 3a). Two sequences recognizing the mouse kcnmb4 open reading frame (see Methods), were inserted into a viral vector to produce adeno associated virus (AAV) (Fig. 3a). Mice were injected with a mix of the two AAVs at postnatal day 18 and two weeks later we performed 2P uncaging of caged glutamate at two individual spines separately and then together near-simultaneously as described above. Analyzing each experiment individually, we found that expression of β4 subunit shRNAs rescued the subthreshold linear integration of synaptic inputs in the basal dendrites of *Fmr1*-KO L5 pyramidal neurons whereas the expression of a nonspecific scrambled version of the shRNA did not (peak linearity: 106.64 ± 2.62 versus 86.61 ± 4.51%, P = 0.001; integral linearity: 103.37 ± 2.88 versus 85.93 ± 5.17%, P = 0.0082; gain linearity: 1.13 ± 0.06 versus 0.87 ± 0.05, P = 0.0063; Fig. 3b, c, d). Similar results were found when the average per mouse was instead considered (peak linearity: 105.36 ± 4.47 versus 88.02 ± 4.68%, P = 0.03; integral linearity: 106.28 ± 3.66 versus 87.67 ± 4.63%, P = 0.0017; gain linearity: 1.20 ± 0.09 versus 0.89 ± 0.06, P = 0.0017; Fig. 3e).

**Figure 3:**
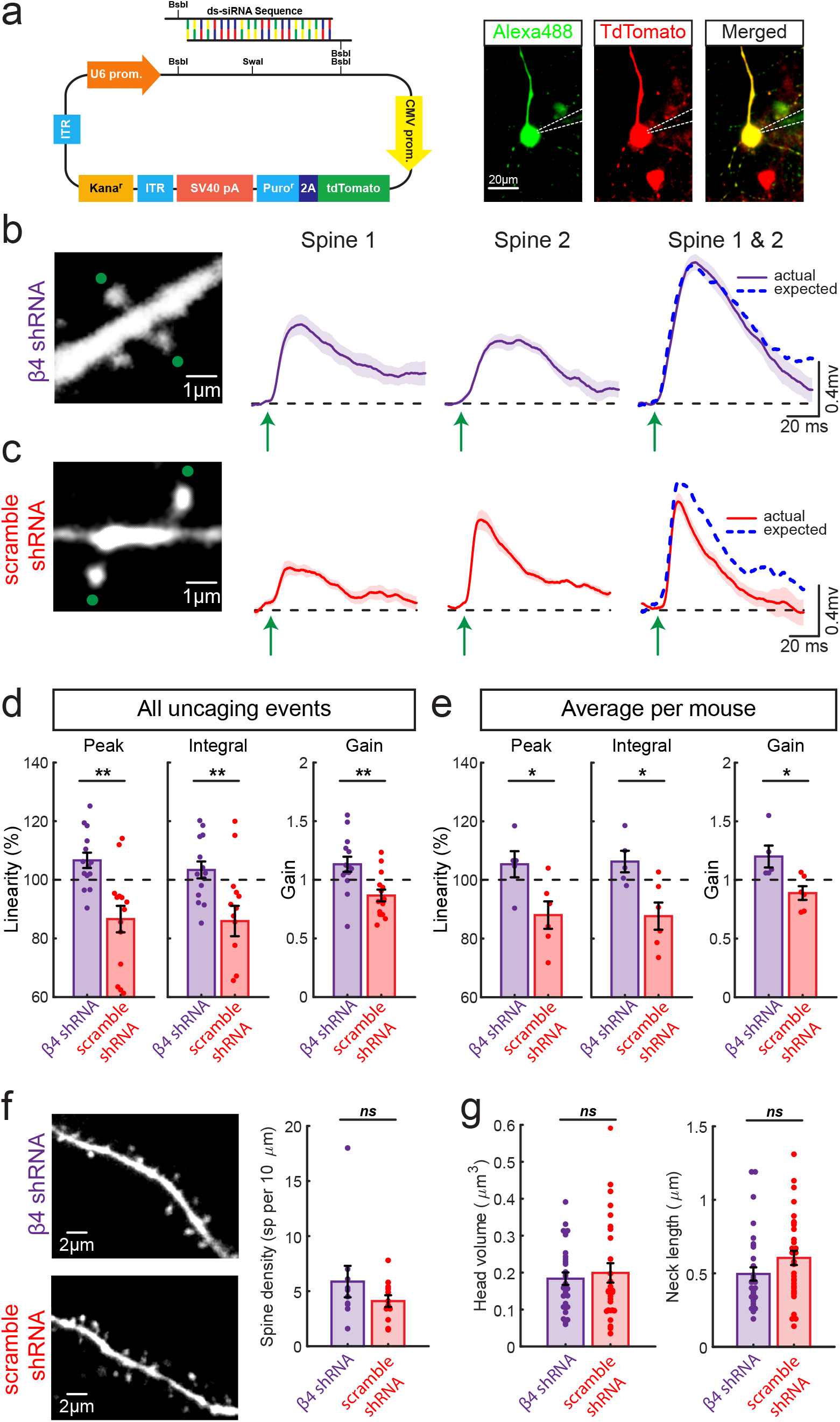
Knock-down of β4 subunit of BK channels rescues sublinearity of synaptic inputs in the basal dendrites of Fmr1-KO L5 pyramidal neurons. (**a**) Schematic of the shRNA vector used to knockdown β4. (**b-c**) shRNA knock-down of the β4 subunit of BK channels rescues the sublinearity in the basal dendrites of *Fmr1*-KO L5 pyramidal neurons (**b**) while injection of scrambled shRNA has no effect (**c**). (**d**) Observed response were not significantly different in amplitude, integral and gain than that expected based on the sum of individual responses in *Fmr1*-KO mice injected with the β4 shRNA, while those injected with a scambled version, the responses summate sublinearly. (**e**) As in **d**, but values were averaged per mouse. (**f-g**)Spine density (**f**) and spine morphology (**g**) are not significantly different in *Fmr1*-KO L5 pyramidal neurons injected with shRNA targeted to the β4 subunit of BK channels versus nonspecific scrambled shRNA in L5 of the visual cortex.

Since activity-dependent spine morphological changes (spine head: (Matsuzaki et al., 2004), neck: (Araya et al., 2014; Tazerart et al., 2020) or both: (Tonnesen et al., 2014)) have been correlated with synaptic efficacy by altering the biochemical and electrical spine properties (reviewed in: Araya, 2014), we analyzed spine shape in mice transfected with either the β4 subunit shRNA or the scramble shRNA. We found that spine head volume and neck length, as well as spine density remained the same in the basal dendrites of *Fmr1*-KO L5 pyramidal neurons transduced with shRNA against the β4 subunit of BK channels versus the scrambled version (spine density: 5.6 ± 1.4, n = 10 images versus 4.1 ± 0.5 spines per 10 μm, n = 12 images; P = 0.432; spine head volume: 0.18 ± 0.02, n = 30 spines, versus 0.19 ± 0.03 μm^3^, n = 28 spines, P = 0.932; neck length: 0.50 ± 0.05, n = 33 spines, versus 0.60 ± 0.05 μm, n = 34 spines, P = 0.096, Mann–Whitney test). These findings provide evidence that the altered synaptic integration arises from an increased activity of the regulatory β4 subunit. This increase can be explained by the lack of FRMP-mediated β4 sequestering in *Frm1*-KO mice (Deng et al., 2013) leading to an elevated binding of the β4 subunit to BK channels impacting the integration of incoming synaptic inputs in the basal dendrites L5 pyramidal neurons.

### Biophysical simulations of synaptic integration in proximal dendrites of *Fmr1*-KO and WT L5 pyramidal neurons

To corroborate our experimental observations on the impact that the interaction of β4 subunit with BK channels in spines on the integration of subthreshold excitatory inputs in the basal dendrites of L5 pyramidal neurons, and gain detailed information on the mechanistic underpinings of this process, we turned to multicompartmental simulations in the NEURON environment (Hines & Carnevale, 1997). Briefly, a morphologically realistic L5 pyramidal neuron was built (adapted from Nevian et al., 2007; Tazerart et al., 2022) where two spines were simulated with neck lengths of 1μm and head volumes of 0.18 μm^3^, which matched the morphology of spines probed in our 2P experiments (Fig. 4a). The dendritic spines were connected to a randomly selected basal dendrite located 170 μm away from the soma of the modeled L5 pyramidal neuron. AMPA-receptors(R), NMDA-R, voltage-gated calcium channels, sodium channels and BK channels were placed in the plasma membrane of each spine head compartment (see Methods; Fig. 4a and b). In one set of simulations, a WT L5 pyramidal neuron was modeled by incorporated the activation/deactivation parameters and kinetics of the α-subunit of BK channels, while in another group a *Fmr1*-KO L5 pyramidal neuron was modeled by using those obtained when the β4 subunit is bound to the α-subunit of BK channels(Jaffe et al., 2011). We then simulated our 2P experiments by modeling synaptic inputs onto two individual spines separately and then together (right panels in Fig. 4a,b). The simulations show that synaptic integration occurs linearly in the simulated WT neuron, when α-subunit of BK channels in spines are modeled (Fig. 4a), while inputs integrate sublinearly in the simulated *Fmr1*-KO neuron when the β4 subunit are incorporated in spines (Fig. 4b), emulating our experimental observations. We quantified these results by calculating linearity indices, using either the peak or integral of the EPSP as well as a gain measure (Fig. 4c). In analyzing how many BK channels per spine are open when one versus two spines are activated in WT and *Fmr1*-KO L5 pyramidal neurons, more BK channels are open when 2 spines are activated. However, in the presence of the β4 subunit there is a significantly larger boosts in the number of open BK channels per spine, leading to the sublinear integration of subthreshold excitatory inputs in the basal dendrites of *Fmr1*-KOL5 pyramidal neurons. (Fig. 4d).

**Figure 4:**
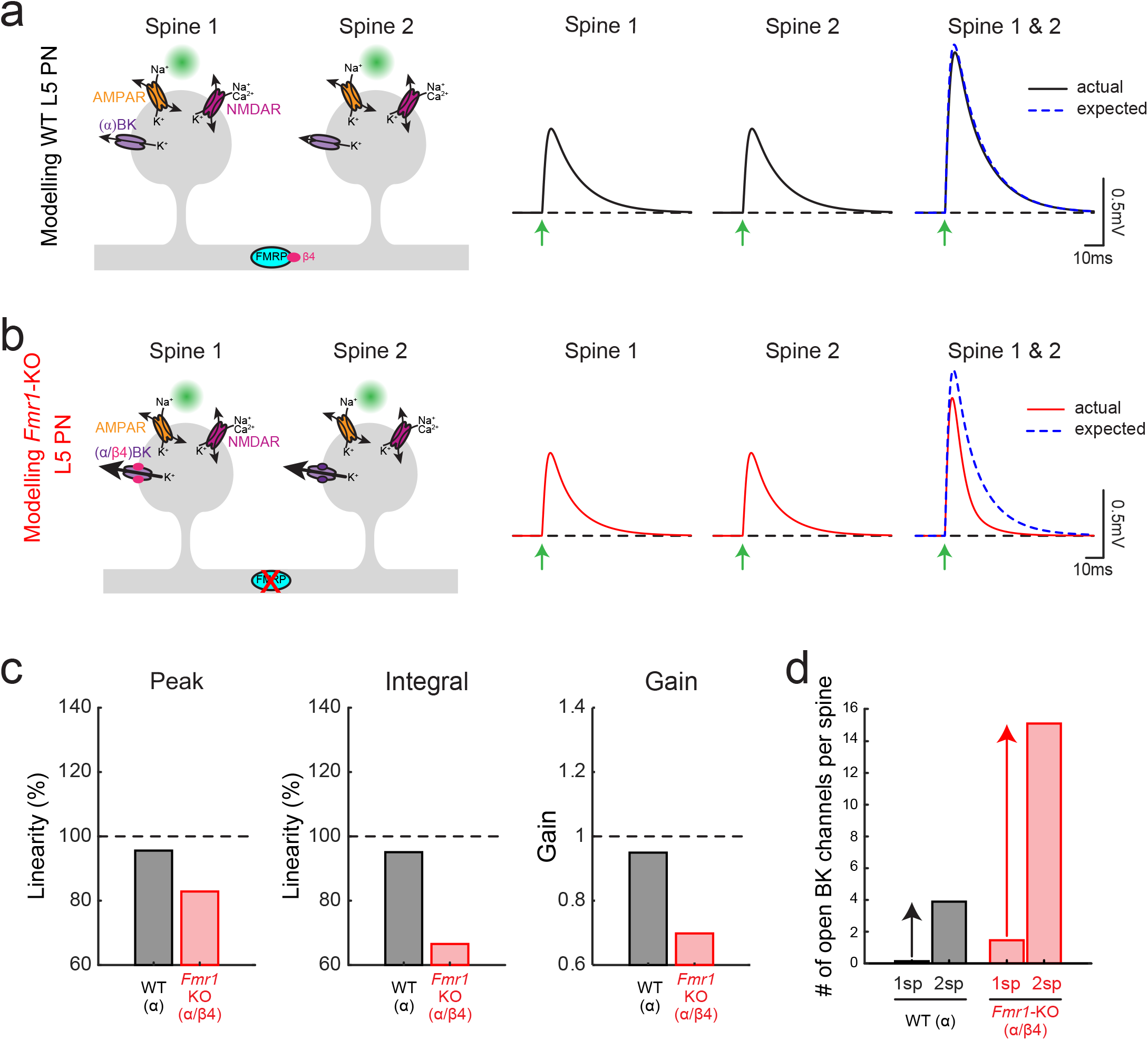
Biophysical modeling shows the effect of the BK channel β4 subunit on synaptic integration in dendritic spines. (**a-b**) *Left panels*: Schematic of the model used to assess synaptic integration in the basal dendrites of a WT L5 pyramidal neuron (**a**) expressing FMRP, which sequesters the β4 subunit so only the α subunit is present in the spine and a *Fmr1*-KO L5 pyramidal neuron (**b**), which does not express FMRP so the β4 subunit is free to bind the BK channel. *Right panels*: Spines were first activated individually (Spine 1 or Spine 2) and then together (Spine 1 & Spine 2) and the EPSP generated at the soma was recorded. Dashed blue traces correspond to the linear sum of individual events of each spine. In the WT neuron, the actual and expected EPSP responses match (compare black and dashed blue traces), whereas in the KO neuron, the actual EPSP response is much less than expected (compare red and dashed blue traces). (**c**) Observed response has smaller amplitude, integral and gain than expected based on the sum of individual responses in *Fmr1*-KO neuron, while in WT mice, the responses summate linearly. (**d**), Number of BK channels open during activation of 1 spine or 2 spines in the modeled WT and *Fmr1*-KO L5 pyramidal neuron.

## DISCUSSION

Previous work has provided evidence that FXS and ASD are characterized by a hyperexcitable neocortex (Bureau et al., 2008; Gibson et al., 2008; Rotschafer and Razak, 2013; Zhang et al., 2014; Booker et al., 2019), thought to be the main contributor to hypersensitivity of sensory stimuli observed in patients (reviewed in: Liu et al., 2021). Our experimental findings presented here challenge this view as they show that the integration of subthreshold excitatory inputs onto the basal dendrites, the main recipients of sensory inputs, of L5 pyramidal neurons in *Fmr1*-KO mouse is sublinear. We further studied the mechanism for this sublinear integration. BK channels and their β4 subunit are localized in the spines of the basal dendrites of L5 pyramidal neurons from WT and *Frm1*-KO mice (Fig. 2) and FMRP (absent in *Fmr1*-KO mice) binds the β4 subunit in the neocortex of WT mice (Deng et al., 2013) thus preventing it from binding to BK channels and altering their activation/deactivation parameters and kinetics. We thus investigated how BK channels and their β4 regulatory subunit affect synaptic integration in *Frm1*-KO L5 pyramidal neurons. Indeed, we found that a knockdown of the β4 regulatory subunit in *Frm1-KO* L5 pyramidal neurons rescued linearity from the sublinear integration previously observed in their basal dendrites. Moreover our numerical simulations corroborate these findings as they show that the linear integration of excitatory synaptic inputs occurs only when the kinetics of the α-subunit of BK channels are modeled, while inputs integrate sublinearly when the β4 subunit are incorporated.

Hypersensitivity to sensory stimuli is described as one of the classical symptoms experienced by individuals with FXS and other forms of ASD (reviewed in: Liu et al., 2021). One might think that being hypersensitive to certain stimuli makes you better able to detect it and perform certain behavioural tasks more aptly, which does not actually seem to consistently be the case for autistic individuals or mouse models of ASD. For example, children with ASD exhibit increased detection thresholds during tactile stimulation (Puts et al., 2014). While it has been shown that *Fmr1*-KO mice exhibit an enhanced acoustic startle response at low sound intensities; they actually show a decreased response to high intensity sounds (Chen and Toth, 2001; Nielsen et al., 2002). *Fmr1*-KO mice also showed impaired learning in a visual discrimination task and took longer to reach expert-level performance comparable to WT animals (Goel et al., 2018). Moreover, this study did not find a significant increase in either spontaneous or visually evoked activity in Layer 2/3 pyramidal neurons of *Fmr1*-KO mice. Thus, the results of these behavioural studies are in agreement with those we report here, in that they cannot consistently be explained by hyperexcitability of the cortex to sensory stimuli. Our data show that near simultaneous activation of two spines in the basal dendrites generates a subthreshold depolarization that summates sublinearly in *Fmr1*-KO mice, whereas, in their WT counterparts, this integration occurs in a linear fashion (Fig. 1) (Araya et al., 2006a). The apical dendrites of *Fmr1*-KO pyramidal neurons, which receive contextual information from other brain areas, are hyperexcitable due to an increased impedance caused by the reduction hyperpolarization-activated cyclic nucleotide–gated (HCN) channels in apical dendrites, resulting in an enhanced temporal summation of excitatory input (Zhang et al., 2014). Thus, taken together, the hypersensitivity characterized in FXS is not simply due to an upweighting of sensory inputs at the level of L5 pyramidal neurons, but instead by a hyposensitivity of sensory inputs and a hypersensitivity of predictive inputs. Specifically, the overrepresentation of predictive inputs onto the apical dendrites of L5 pyramidal neurons, coinciding with a downrepresentation of sensory inputs at the level of the basal dendrites of cortical pyramidal neurons, may be at the core of the behavioral phenotypes associated with FXS. These changes would likely have significant effects on the input/output properties of L5 pyramidal neurons, and in the ability of the cortex to decipher sensory signals in order to make predictions about the future. Moreover, the results from the present study highlight the need to study cortical feedforward and feedback pathways independently, their associations and cortical pyramidal neuron’s output in FXS and other forms of ASD.

Although the present study focused on L5 pyramidal neurons in *Fmr1*-KO mice, there have been many studies on how FXS and ASD in general alters the activity of cortical interneuron. Many mouse models of ASD exhibit altered interneuron function and development (Lunden et al., 2019). Specifically, compared to WT animals, *Fmr1*-KO mice displayed delayed learning of a visual discrimination task, and reduced activity of parvalbumin (PV) interneurons in the visual cortex. Restoring PV cell activity accelerated the learning in *Fmr1*-KO mice (Goel et al., 2018). Moreover, the somatosensory cortex of Fmr1-KO mice have significant reduction the density of PV (Selby et al., 2007). Thus, it is likely that the stereotypical behaviour observed in FXS is caused by complex alteration in the cortical circuits, including both excitatory and inhibitory neurons.

Studies investigating how the BK channel β4 subunit impacts the activity of hippocampal and neocortical pyramidal neurons have shown that FMRP interacts with the β4 subunit and that the lack of FMRP leads to excessive broadening of pyramidal neuron action potentials, and consequently their inter-spike-interval during repetitive activity (Deng et al., 2013). Thus, FMRP can regulate neural firing patterns through a BK channel-mediated mechanism by the interaction of FMRP and the BK channel β4 subunits (Deng et al., 2013). The effect of the β4 subunit on BK channels is complex as it decreases the probability of BK channels opening at low intracellular calcium concentrations but promotes channel opening at high intracellular calcium concentrations while also slowing down activation and deactivation kinetics (Brenner et al., 2000; Torres et al., 2007). Thus, how the β4 subunit will affect BK channel opening depends on intracellular calcium levels. In the present study, we focused on how the BK channel β4 subunit influences the integration of subthreshold inputs (i.e., uEPSPs). Taken together, our results along with previous studies show that in *Fmr1*-KO pyramidal neurons, the β4-bound BK channels causes (1) subthreshold inputs to be integrated sublinearly in the basal dendrites and (2) alterations in the timing of action potentials and information transmission compared to WT mice. Both of these defects can severely influence how sensory information is transmitted to downstream neurons from L5 pyramidal neurons.

The findings presented here are quite remarkable since, to our knowledge, this is the first demonstration of hyposensitivity of sensory inputs at the level of L5 pyramidal neurons in a mouse model of ASD. Specifically, these results reveal a hyposensitivity of basal integration of sensory inputs in *Fmr1*-KO L5 pyramidal neurons, contradicting the conventional view that FXS is characterized by sensory hypersensitivity at the cellular level, but providing instead a more complete picture where the differential integration of feedforward and feedback inputs in L5 pyramidal neurons could be at the core of the cellular basis of FXS.

## METHODS

### Animals

C57B/6 (WT, RRID:IMSR_JAX: 000664) and *Fmr1*-KO (B6.129P2-*Fmr1^tm1Cgr^*/J, RRID:IMSR_JAX:003025) and *Fmr1*-KO;thy1GFPM (*Fmr1*-KO backcrossed to Tg(Thy1-EGFP)MJrs/J, RRID:IMSR_JAX:007788) mice were used in this study and housed on a 12-h light/dark cycle with ambient temperature 20–24 °C and 40–70% humidity.

### Brain slice preparation and electrophysiology

Mice (P14-40), anesthetized with isoflurane, were decapitated, and their brains dissected and placed in cold (4°C) carbogenated sucrose cutting solution containing (in mM) 27 NaHCO_3_, 1.5 NaH_2_PO_4_, 222 sucrose, 2.6 KCl, 1 CaCl_2_, and 3 MgSO_4_. Coronal brain slices (300-μm-thick) of visual cortex were prepared using a Vibratome (VT1000 S, Leica, Germany) and slices were incubated for 30min at 32°C in ACSF (in mM: 126 NaCl, 26 NaHCO_3_, 10 dextrose, 1.15 NaH_2_PO_4_, 3 KCl, 2 CaCl_2_, 2 MgSO_4_) and then at room temperature until ready for use. MultiClamp 700 B amplifiers (Molecular Devices) were used for electrophysiological recordings of L5 pyramidal neurons with a patch electrode (4-7MΩ) filled with internal solution containing (in mM): 0.1 Alexa-568, 130 Potassium D-Gluconic Acid (Potassium Gluconate), 2 MgCl_2_, 5 KCl, 10 HEPES, 2 MgATP, 0.3 NaGTP, pH 7.4, and 0.4% Biocytin. DIC optics were used to clearly visualize and patch the soma of L5 pyramidal neurons.

### Two-photon imaging and two-photon uncaging of glutamate

Two-photon imaging was performed using a custom-built two-photon laser scanning microscope (Mitchell et al., 2019; Tazerart et al., 2020), consisting of (1) a Prairie scan head (Bruker) mounted on an Olympus BX51WI microscope with a ×60, 0.9 NA water-immersion objective; (2) a tunable Ti-Sapphire laser (Chameleon Ultra-II, Coherent, >3 W, 140-fs pulses, 80 MHz repetition rate), (3) two photomultiplier tubes (PMTs) for fluorescence detection. Fluorescence images were detected with Prairie View 5.4 software (Bruker).

Fifteen minutes following break-in, two-photon scanning images of basal dendrites were obtained with 720 nm at low power (< 5 mW on sample) excitation light and collected with a PMT. Two-photon uncaging of 4-methoxy-7-nitroindolinyl (MNI)-caged L-glutamate (2.5 mM; Tocris) was performed using a 4 ms pulse at 720 nm and ~30mW on sample (Araya et al., 2006a) with the uncaging spot ~0.3 μm away from the upper edge of the selected spine head. The uEPSPs were recorded with the patch pipette at the soma of L5 pyramidal neurons. To assess the integration of individual synaptic inputs onto the basal dendrites of L5 pyramidal neurons, two-photon uncaging of MNI-glutamate was used. Uncaging was performed first at two neighbouring spines separately and then together (interstimulus interval of <0.1 ms) with a 2s delay in between each uncaging event. This sequence was repeated 10 times. Recordings were obtained using a MultiClamp 700B amplifier (Axon Instruments) interfaced to a dedicated computer by a BNC-2090A data acquisition board (National Instruments). The electrophysiological signals were acquired at 10 kHz using the *PackIO* open-source software package (*http://www.packio.org*).

### Data analysis

Offline data analysis was performed with the MATLAB (Mathworks) *EphysViewer* package (*https://github.com/apacker83/EphysViewer*) and custom written algorithms. We first quantified our results by calculating linearity indices, using either the peak or integral of the uEPSP, as described in the following equations:

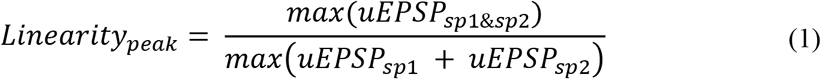

and

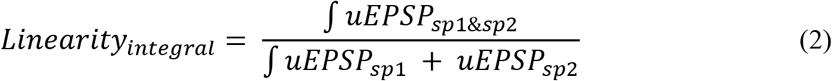

where *uEPSP_sp1_, uEPSP_sp2_* and *uEPSP_sp1 & sp2_* is the time varying uEPSP response when we applied 2P uncaging of glutamate at spine 1, spine 2 and then together near-simultaneously, respectively.

Linear optimization techniques were also used to quantify the linearity of synaptic inputs for WT and *Fmr1*-KO mice. Specifically, the uEPSP response to near-simultaneous uncaging of neighbouring spines was modeled using the following equation:

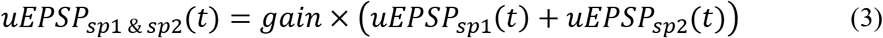

where *uEPSP_sp1_*(*t*), *uEPSP_sp2_*(*t*) and *uEPSP_sp1 & sp2_*(*t*) is the time varying uEPSP response when we applied 2P uncaging of glutamate at spine 1, spine 2 and then together near-simultaneously, respectively and *gain* is measure of how close the expected (based on the arithmetic sum) and actual responses to uncaging spine 1 and spine 2 are to each other – a measure of linearity of the time-varying uEPSP response.

Analysis of the spine morphology was performed in the open source image processing package Fiji (NIH). Specifically, the spine neck was measured as the proximal edge of the spine head to the edge of the dendrite. In cases where the spine topology could not be precisely determined, the spine neck length was estimated as the shortest orthogonal distance between the base of the spine head and the edge of the dendrite. When the spine neck could not be measured, a minimum value of 0.2 μm was used. For the spine head size, the longest and corresponding orthogonal diameter was measured. These measures were computed from a Gaussian curve fit to the fluorescent profile of these axes, generated from a z-stack over the entire spine heads (Δz = 0.4 μm). The corresponding spine head volume (V, in μm^3^) was estimated using the following formula:

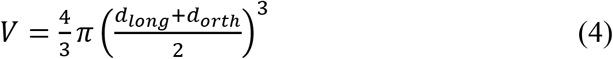

where d_long_ and d_orth_ are the FWHM of the spine head along the longest and corresponding orthogonal dimension, respectively. The FWHM was calculated using the standard deviation (σ) of the Gaussian curve fit to the fluorescent profile of the spine heads (FWHM = 2.335 σ).

#### Synaptoneurosome (SN) preparation

An adaptation of a previously described method was used (Villasana et al., 2006; Seibt et al., 2012) as follows: P14 mouse visual cortex was dissected, flash frozen and stored at - 80°C. Individual samples (approx. 1 mg) were homogenized in 700 μL SN lysis buffer (10 mM Hepes, pH 7.4, containing proteinase inhibitor) using a 2 mL Kimble-Chase tissue grinder (Thermo Fisher K885300-0002) and applying 6 loose pestle and 12 tight pestle strokes. A 150 μL aliquot of this total protein (TP) fraction was boiled in 10% SDS for 10 min and stored. The remaining fraction was centrifuged at 2000xg for 2 min at 4°C to remove cellular and nuclear debris. The supernatant was loaded into a 1 mL syringe and filtered through three layers of pre-wetted 100 μm pore nylon filter (Millipore NY1H04700). The filtrate was directly loaded into 5-μm-pore centrifugal filters (Ultrafree-CL, Millipore UFC40SV25) and centrifuged at 5000xg for 15 min in a fixed angle rotor at 4 °C. The supernatant was carefully removed, and the loose pellet resuspended in boiling SN buffer (containing 2% SDS), boiled for 10 min and store at – 80°C (SN protein fraction). Protein concentration was determined with the BCA protein assay kit (Thermo Fisher 23227) using a bovine serum albumin (BSA) standard curve.

#### Western Blot

An adaptation of a previously described method was used (Miranda-Rottmann et al., 2010) as follows: protein samples were separated in NuPAGE™ 3-8% Tris-Acetate Protein Gels (Thermo Fisher EA03755) in Tris-Acetate-EDTA running buffer (Thermo Fisher BP13354) at 200V for 1h for αBK and Bis-Tris 4-12% polyacrylamide gradient gels (Thermo Fisher NP0323) in MES-SDS running buffer (Thermo Fisher B0002) at 200V for 25 min for β4 subunit. Separated proteins were transferred into a PVDV membrane using a mini trans-blot cell (BioRad 170-3930) at 200V for 1h. Ponceau red staining was performed after transfer and recorded. Membranes were blocked in 0.1% Tween20 in Phosphate buffered saline (PBS) starting block (Thermo Fisher 37538) and incubated with primary antibodies diluted in the same buffer: anti-BK channel 1:500 (Antibodies Inc 75-022) or 1:100 anti-BKbeta4 K+ channel (Antibodies Inc 75-086) and after striping a mix of anti-γ-tubulin 1:10,000 (Sigma-Aldrich,T5326,) and anti-PSD95 1:5,000 (Antibodies Inc 75-028). Signal was developed using ECL substrate (BioRad 170-5060) and recorded on film. Densitometric analysis was performed with the open-source program FIJI (Schindelin et al., 2012) and data was plotted using Prism9 (GraphPad)

#### Immunofluorescence

Anesthetized P17-27 Fmr1-KO;thy1GFPM mice were intracardially perfused with 4% PFA in PBS. Post-fixing of the brain was done in 4% PFA for 2h at 4° followed by washes in PBS and dehydrated in 30% sucrose at 4°C and then mounted in OCT by freezing in a bath of 2-methylbutane at −50°C. The 40 μm cryosections were cut at −18°C and placed over glass slides treated with tissue capture pen (Electron Microscopy Sciences 71314) and stored at −80°C. Cryosections were permeabilized and blocked in TSA blocking reagent (Perkin Elmer FP1020). Primary antibodies used were anti-BK 1:100 (Antibodies Inc AB_224953), 1:100 anti-BKbeta4 K+ channel (Antibodies Inc 75-086) and anti-GFP 1:500 (Rockland 600-101-215). Incubations were performed for 14h at 4° C followed by incubation with secondary antibodies: Alexa 568 anti-mouse 1:200 (Thermo Fisher A10037), and Alexa 488 anti-goat 1:200 (Thermo Fisher A11055) for 2h at room temperature (RT) in TSA blocking reagent. Washes were done in PBS. The samples were mounted in ProLong Diamond Antifade Mountant with DAPI (Thermo Fisher P36971) or Vectashield (Vector Labs H-1200) and the basal dendrites from L5 pyramidal neurons were imaged using a Zeiss LSM700 confocal microscope (IRIC bio-imaging core).

### Virus injection

To perform the knockdown of the β4 subunit of BK channels, a mix of two different adeno associated virus serotype 1 (AAV1) were used, each containing a different shRNA based on kcnmb4 the mouse sequence (NM_021452): shkcnb4-NM_021452.126s1c1: 5’-CCGGCCCGCCUUGCAGGAUCUGCAACUCGAGUUGCAGAUCCUGCAAGGCGGGUUUUUG-3’ (binding nucleotides 126-146) and shkcnb4_NM_021452.1-266s1c: 5’-CCGGCGUGAACAACUCCGAGUCCAACUCGAGUUGGACUCGGAGUUGUUCACGUUUUUG-3’ (binding nucleotides 266-189) or a control custom scramble shRNA: 5’-CCGGACCACCGTCGAATCGTGACAACTCGAGTTGTCACGATTCGACGGTGGT TTTTTG-3’ cloned downstream of the U6 promoter in a pAAV-shRNA-tdTomato vector and prepared at a concentration of 10×10^13^ GC/mL (abm, Canada). The morning of the surgery, *Fmr1*-KO mice (P18) were given Sustained-Release Buprenorphine (1mg/kg). Mice were then deeply anesthetized with 3% isoflurane, confirmed by the absence of a toe pinch response and placed into a stereotaxic frame (Kopf Instruments, California). An incision was made into the disinfected bare skin to expose lambda and bregma landmarks. A small hole was drilled into each side of the skull at 3mm posterior and 2mm lateral to bregma. The virus was loaded into a Hamilton syringe and injected 0.5 mm below the pial surface. The virus (450 μL) was injected bilaterally, over the course of 20 min followed by a 5 min wait period before the syringe was slowly and carefully removed. The incision was sutured and mice were returned to their home cage and given 2 weeks to recover and obtain a good virus expression. 2P uncaging experiments, as described above, were performed on L5 pyramidal neurons expressing the reporter gene tdTomato.

### Modelling

We used a morphologically realistic L5 pyramidal neuron NEURON model (Nevian et al., 2007; Tazerart et al., JPhysiol 2022) to study the effects of the β4 subunit on synaptic integration. Two spines, with neck lengths of 1 μm and spine head volumes of 0.18 μm^3^, were inserted 170 μm from the soma on a randomly selected basal dendrite. The neck width was estimated based on the head volume using previously reported linear relationships that found a positive correlation between head size and neck width in distal dendrites (Arellano et al., 2007), also been observed *in-vivo* using STED nanoscopy (Steffens et al., 2021) and other cortical pyramidal neurons (Ofer et al., 2021). Neck and spine head axial resistances were set to 500 and 150 MΩ respectively based on published values (O’Donnell et al., 2011; Harnett et al., 2012). High voltage-dependent Ca^2+^ channels (HVA, https://senselab.med.yale.edu/modeldb/ShowModel?model=135787&file=/ShuEtAl20062007/ca.mod#tabs-2), Ca^2+^ buffering/removal mechanisms (resting concentration of 1nM (Destexhe et al., 1993)), Nav1.6 sodium channels (3 S/cm^2^; (Mercer et al., 2007)) and BK channels (14 S/cm^2^) (Jaffe et al., 2011; Tazerart et al., 2022) were inserted in the spine heads. In one set of simulations, we incorporated the activation/deactivation parameters and kinetics of the α-subunit of BK channels, while in another set of simulations we used those obtained when the β4 subunit is bound to BK channels (Jaffe et al., 2011). To activate individual synapses, we used an ionotropic glutamate receptor stimulation mechanism (Polsky et al., 2009) such that AMPA and NMDA conductances (10 nS and 0.086 nS, respectively) were calibrated to obtain physiologically realistic EPSPs at the soma (~0.5-0.6 mV).

### Ethics

These studies were performed in compliance with experimental protocols (13-185, 15-002, 16-011, 17-012, 18-011, and 19-018) approved by the Comité de déontologie de l’expérimentation sur les animaux (CDEA) of the University of Montreal and protocol 2020-2634 approved by the Comité institutionnel des bonnes pratiques animales en recherche (CIBPAR) of the Centre de Recherche, CHU Ste-Justine.

## Figure Legends

**Supplementary Figure 1:**
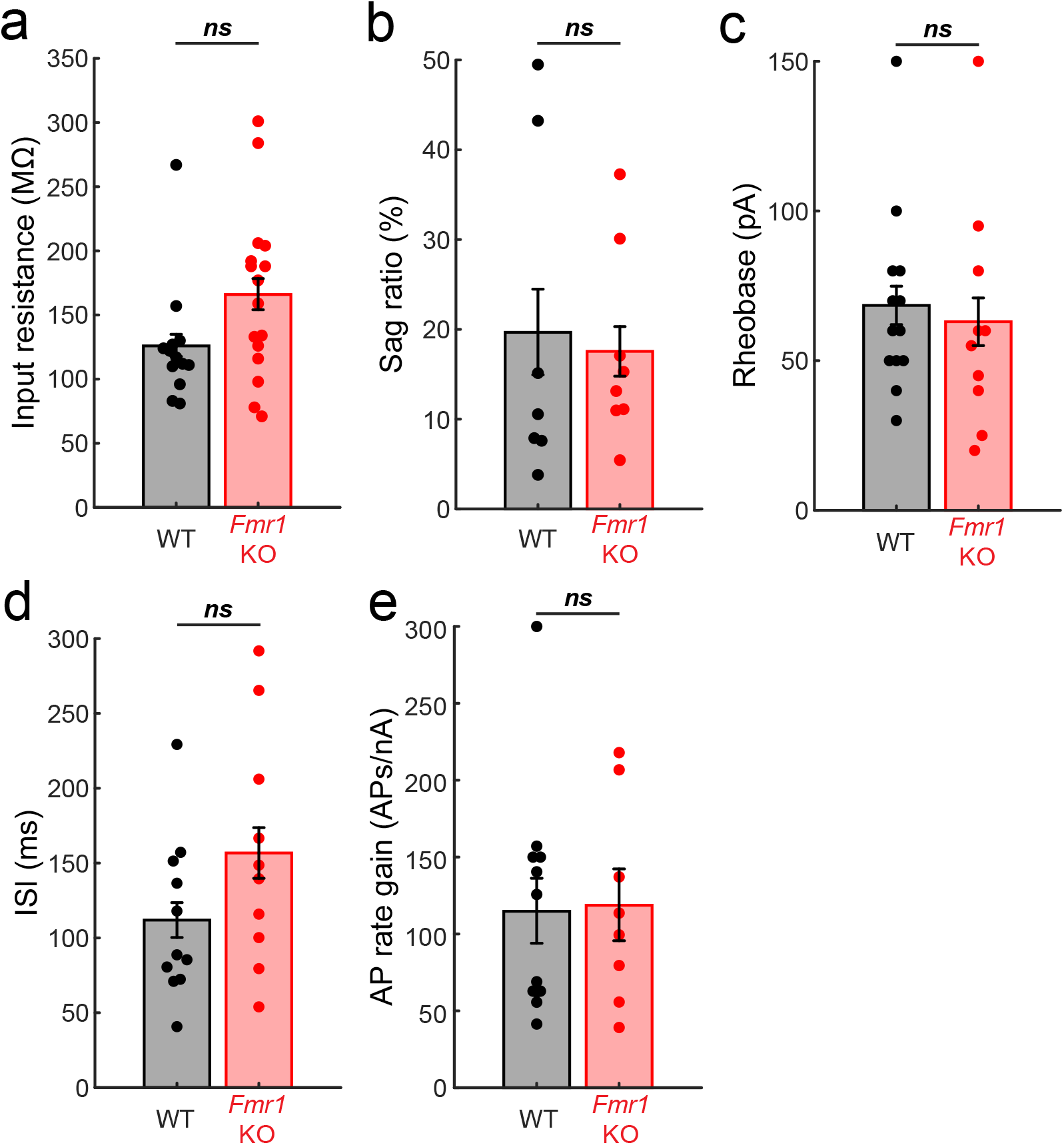
Intrinsic cellular properties of *Fmr1*-KO and WT L5 pyramidal neurons. No significant difference in input resistance (**a**), sag ratio (**b**), rheobase (**c**), interspike interval (**d**) and action potential rate gain as a function of current injected (**e**) between *Fmr1*-KO and WT L5 pyramidal neurons.

**Supplementary Figure 2:**
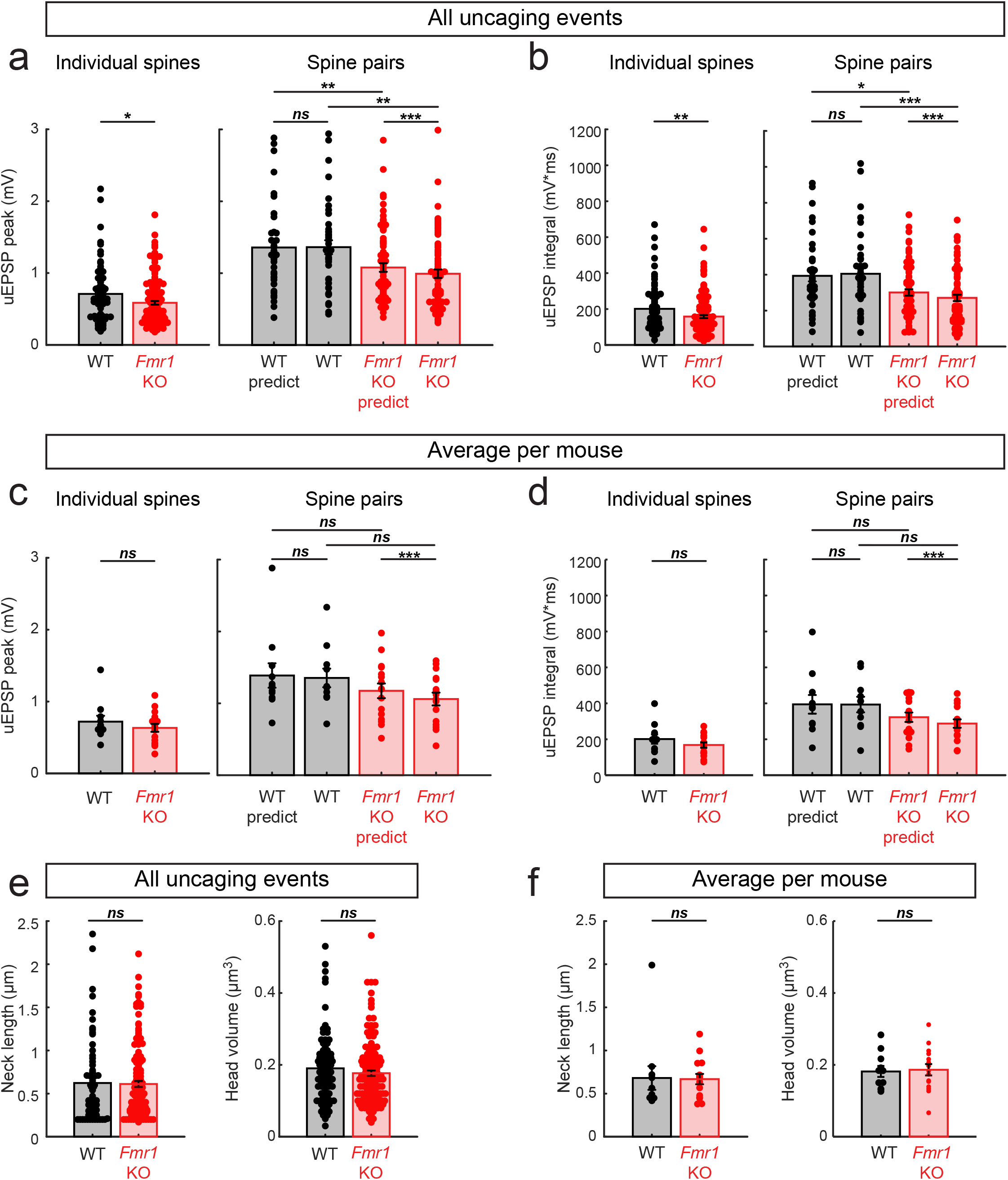
Raw uEPSP amplitudes and integral and spine morphology. (**a**) Raw uEPSP peak values for all experiments shown in Fig. 1. (**b**) Raw uEPSP integral values for all experiments shown in Fig. 1. (**c**) As in **a**, but uEPSP peak values were averaged per mouse. (**d**) As in **b**, but uEPSP integral values were averaged per mouse. (**e**) Neck length and head volume for all spines activated during experiments presented in Fig. 1. (**f**) As in **e**, but values were averaged per mouse.

**Supplementary Figure 3:**
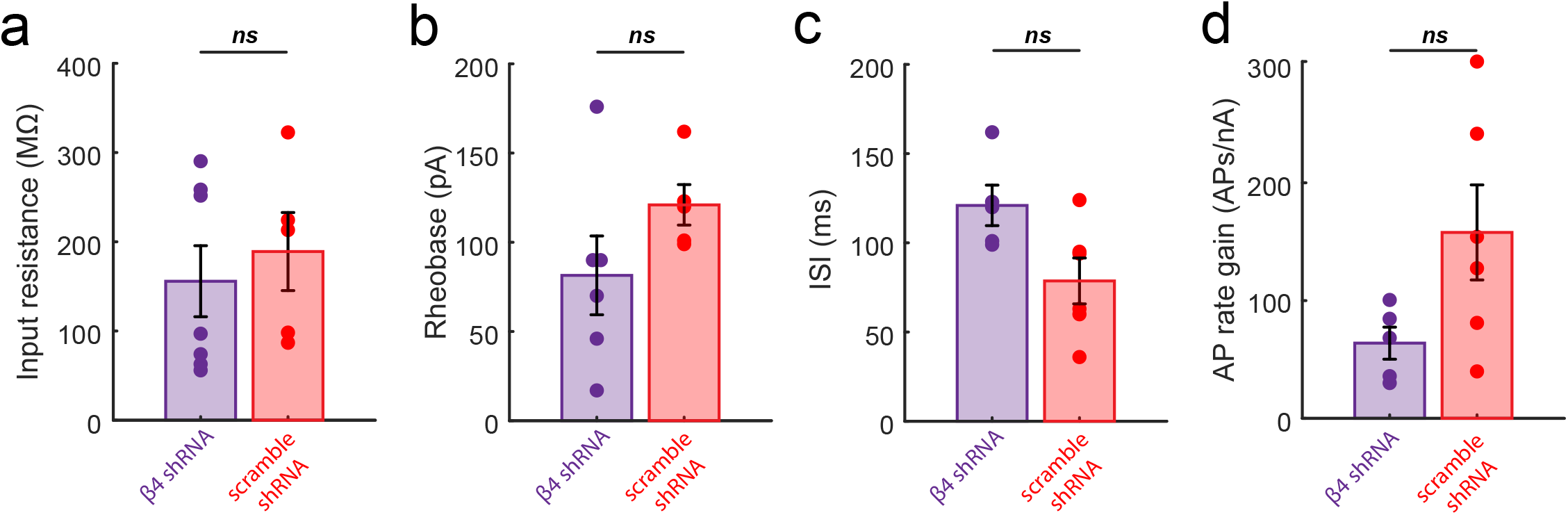
Intrinsic cellular properties of *Fmr1*-KO L5 pyramidal neurons following injections of shRNA against BK channel β4 subunit and scrambled version. No significant difference in input resistance (**a**), rheobase (**b**), interspike interval (**c**) and action potential rate gain as a function of current injected (**d**) between *Fmr1*-KO L5 pyramidal neurons injected with shRNA against BK channel β4 subunit and scrambled version.

